# Epigenetic mutational landscape in breast cancer: role of the histone methyltransferase gene *KMT2D* in triple negative tumors

**DOI:** 10.1101/486233

**Authors:** Sara Morcillo García, María del Mar Noblejas-López, Cristina Nieto-Jiménez, Javier Perez-Peña, Miriam Nuncia-Cantarero, Balázs Győrffy, Eitan Amir, Atanasio Pandiella, Eva Galán-Moya, Alberto Ocaña

## Abstract

**Purpose:** Epigenetic regulating proteins like histone methyltransferases produce variations in several functions, some of them associated with the generation of oncogenic processes. Mutations of genes involved in these functions have been recently associated with cancer, and strategies to modulate their activity are currently in clinical development.

**Methods:** By using data extracted from the METABRIC study, we searched for mutated genes linked with detrimental outcome in invasive breast carcinoma (n = 772). Then, we used downstream signatures for each mutated gene to associate that signature with clinical prognosis using the online tool “Genotype-2-Outcome” (http://www.g-2-o.com). Next, we performed functional annotation analyses to classify genes by functions, and focused on those associated with the epigenetic machinery.

**Results:** We identified *KMT2D, SETD1A* and *SETD2*, included in the lysine methyltransferase activity function, as linked with poor prognosis in invasive breast cancer. *KMT2D,* which codes for a histone methyltransferase that acts as a transcriptional regulator, was mutated in 6% of triple negative breast tumors and found to be linked to poor survival. Genes regulated by *KMT2D* included *RAC3, KRT23,* or *KRT14,* among others, which are involved in cell communication and signal transduction. Finally, low expression of *KMT2D* at the transcriptomic level, which mirror what happens when *KMT2D* is mutated and functionally inactive, confirmed its prognostic value.

**Conclusion:** In the present work, we describe epigenetic modulating genes which are found to be mutated in breast cancer. We identify the histone methyltransferase KMT2D, which is mutated in 6% of triple negative tumors and linked with poor survival.

## Introduction

Advances in the analyses of the genomic landscape of human cancers have permitted the identification of different molecular alterations, including mutations, copy number variations, or gene rearrangements, which may be linked with the genesis and maintenance of tumors [1,2]. Unfortunately, for most of the identified molecular alterations, limited druggable opportunities exist [1,2]. Very well-known exceptions include inhibition of protein kinase activity, when that alteration affects a kinase [2]. This has been the case for agents targeting mutated or amplified protein kinases, such as EGFR or HER2 in lung and breast cancers [3–5]. In a similar manner, chromosomal rearrangements can produce fusion proteins, like Trk fusion proteins, with kinase activity amenable for pharmacological inhibition [6,7].

Changes at the genome not directly produced by an alteration of the nucleotide sequence of the DNA are known as epigenetic modifications [8]. Alterations in proteins involved in epigenetic regulation can affect genetic programs that can in turn impact on several cellular functions. Ultimately, such genomic alterations can translate into different diseases, from cancer to neurological alterations or aging disorders, among others [8,9]. Epigenetic regulating proteins include enzymes involved in histone modifications, histone proteins, chromatin remodeling complexes or DNA methylation enzymes [8]. Mutations at genes coding for proteins involved in several of these functions have been already described, and some of them have been associated with cancer [10]. Therefore, inhibition of epigenetic proteins can have a wide effect impacting on the expression of multiple genes, affecting multiple pathways at the same time [10]. In this context, agents that target epigenetic enzymes have been recently described and are currently in clinical development [11].

In this study, we evaluated the mutational status of genes involved in epigenetic control in breast cancer, identifying *KMT2D* as mutated in around 6% of triple negative tumors and linked with a particular detrimental prognosis.

## Material and methods

### Identification of Breast Cancer mutated genes

Data was extracted from the Breast Cancer METABRIC study (n = 2509), contained at cBioPortal (http://www.cbioportal.org). First, we searched for mutated genes in those samples from Invasive Breast Carcinoma patients (n = 772), including luminal A, luminal B, HER2+ and basal-like. Genes that were mutated in more than 2.5% of the patients were identified. The frequency of mutations was independently confirmed using the TCGA database (n = 818).

### Functional analyses

For the functional annotation analysis of the set of mutated genes, the gene list enrichment analyses tool DAVID Bioinformatics Resources 6.8 (https://david.ncifcrf.gov/) was used. To do so, genes with a mutation frequency greater than 2.5% and linked with poor prognosis were selected.

For the functional analysis of the KMT2D-associated gene signature (S1 Table), the online Enrichr tool was used (http://www.amp.pharm.mssm.edu/Enrichr/). An adjusted *p*-value <0.05 was applied to select enriched gene-sets. Genes were separated into overexpressed and underexpressed and “KEGG 2015” option was chosen for the analyses and the calculation of the “combined score”.

### Outcome analyses

To evaluate the relationship between the presence of mutated genes and patient clinical outcome, the Genotype-2-Outcome online tool (http://www.g-2-o.com) [12] was used (S1 table). This publicly available database allows the evaluation of clinical outcome for all breast cancer subtypes (All, Triple Negative Breast Cancer, Luminal A, Luminal B and HER2+) by exploring the association with prognosis of a specific transcriptomic signature associated with that mutation.

To evaluate the relationship between the expression of the genes and patient clinical prognosis, the KM Plotter Online Tool (http://www.kmplot.com) [13,14][13,14][12, 13] was used. This database permits the evaluation of overall survival (OS) and relapse-free survival (RFS) in basal-like, luminal A, luminal B, HER2+ and triple negative breast cancers.

For both outcome analyses, patients were separated according to median values. Patients above the threshold were considered to have a “high” expression while patients below the threshold were defined as those with “low” expression.

### Evaluation of KMT2D mutations

Data contained at cBioportal (http://www.cbioportal.org) was used to identify mutations in *KMT2D.* Mutation Assessor (http://www.mutationassessor.org), SIFT (http://sift.bii.a-star.edu.sg/) and PolyPhen-2 (http://genetics.bwh.harvard.edu/pph2/) databases were used to evaluate the effect of the mutation on *KMT2D* functionality.

## Results

By using the METABRIC database, we identified 172 mutated genes in the analyses of the 772 samples from invasive breast tumors. We found that 59 out of the 172 genes were mutated in more than 2.5% of the samples. Next, we evaluated the impact of these genes on patient outcome using the online tool Genotype-2-Outcome (http://www.G-2-O.com/)[12] (Figure 1A). This application identifies the transcriptomic signature associated with the presence of the mutation in patients. Using this approach, 44 of the mutated genes had an associated signature linked to detrimental prognosis in breast cancer (Figure 1A).

**Figure 1.**
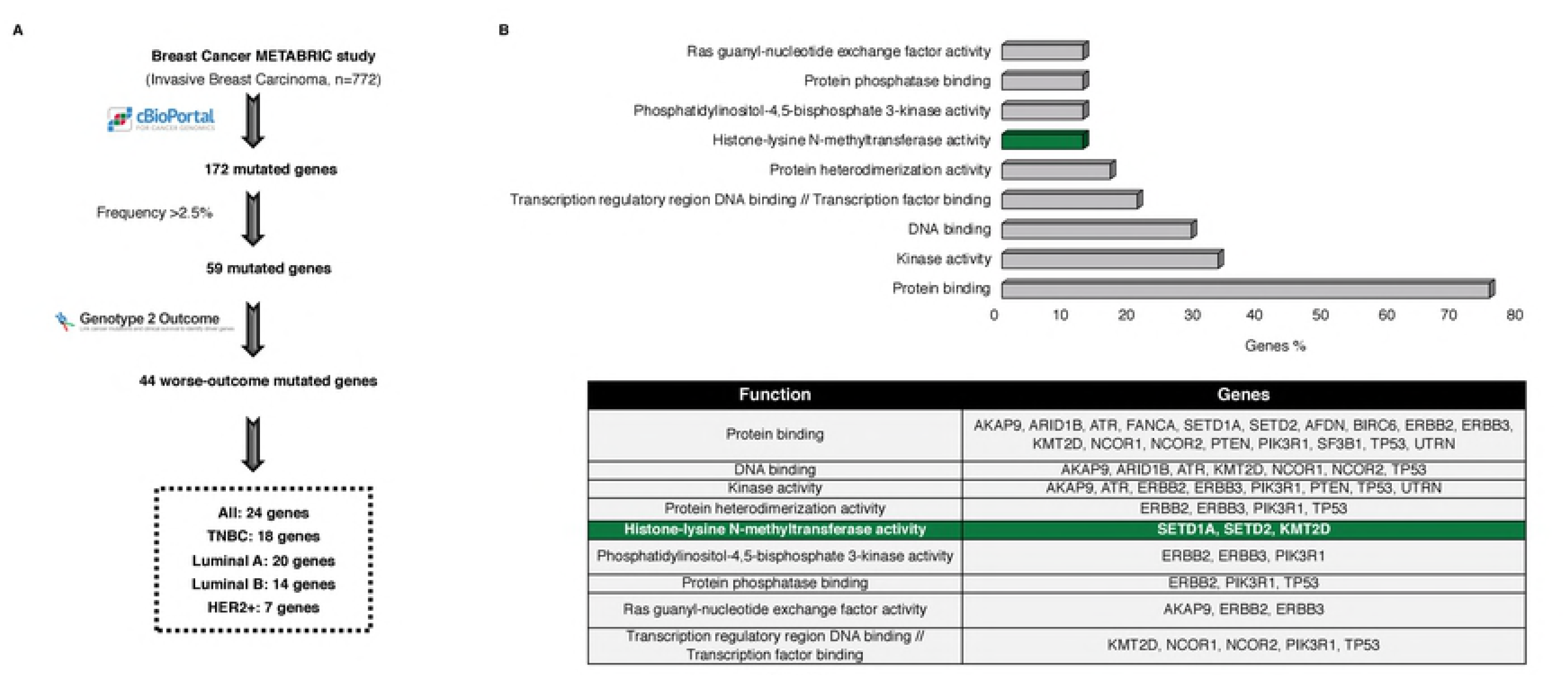
Whole genome mutational profiling and identification of histone-lysine methyltransferase activity as deregulated in breast cancer. A. Flow chart of the study, in which the METABRIC dataset was used to identify breast cancer mutated genes associated with worse outcome. B. Functional analyses of the mutated genes associated with worse outcome, using DAVID Bioinformatics Resources 6.8 tool, and found in more than 2.5% of breast cancer samples analyzed. The table shows the list of the mutated genes contained in each function.

To get insides into the biological function of the mutated genes, we performed a functional annotation analysis. Protein binding, kinase activity, DNA binding and transcription factor binding were among the identified functions which grouped more genes (Figure 1B). Then, the mutational frequency of the identified genes for all breast cancer subtypes was studied. Mutations in some of the genes have been widely described in breast cancer, as is the case for *TP53,* in luminal and HER2+ tumors (Figure 2A). In the case of TNBC, mutated genes displaying higher frequency, more than 8%, included *SYNE1, CDH1* and *DNAH11* (Figure 2A). In HER2+ disease, *PIK3CA* was mutated in more than 40% of tumors. Of note, mutated genes found in TNBC tumors showed a broader range of functions than the other subtypes (Figure 2B).

**Figure 2.**
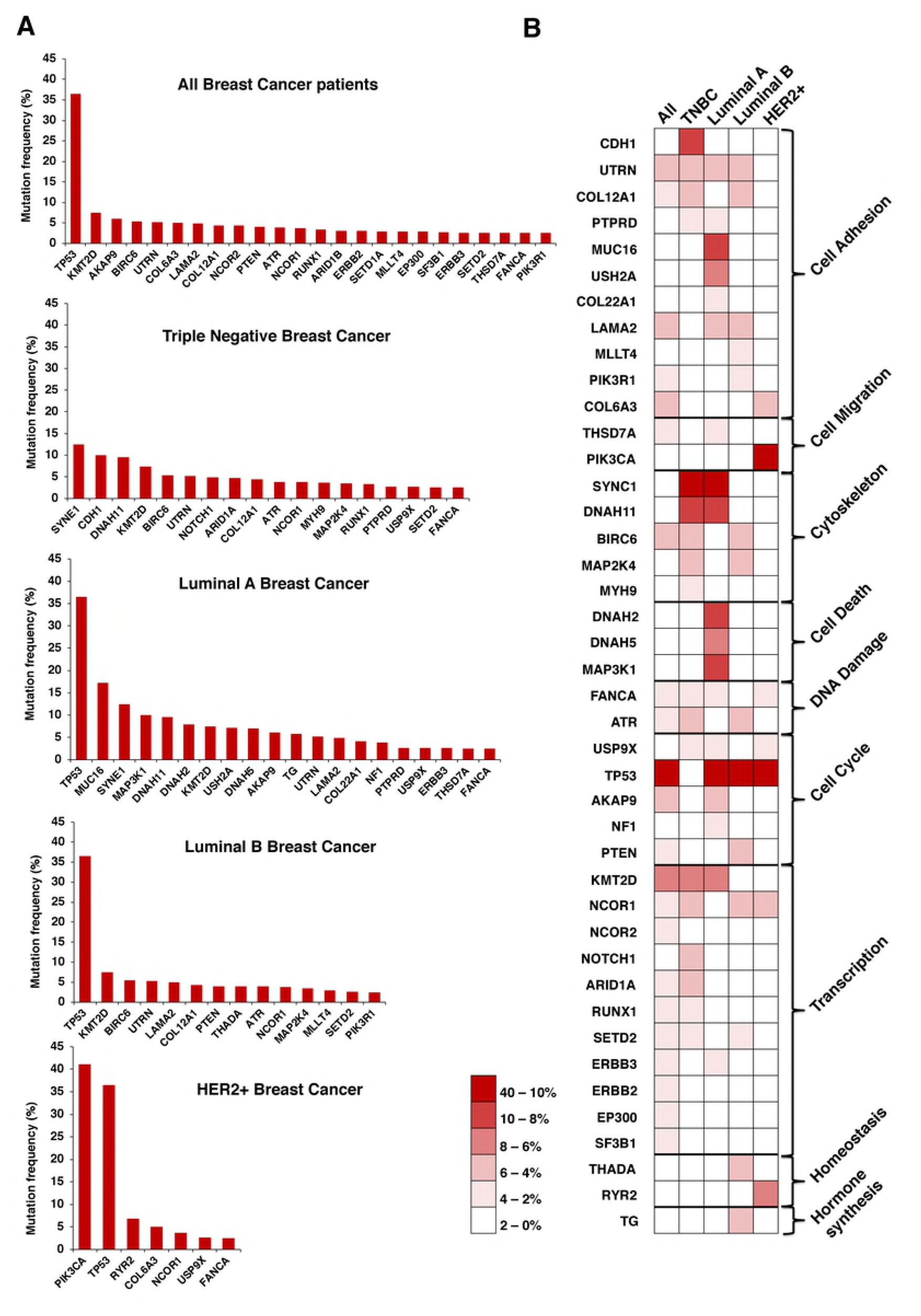
Mutational profile by breast cancer subtype, and association with biological functions. A. Graphs displayed the mutation frequency of those genes mutated in more than 2.5% of patients for all and each breast cancer subtype. B. Heat map of the mutation frequency and the functions of the identified genes for each breast cancer subtypes. The percentage of mutated cases is displayed in the legend.

Because epigenetic enzymes are currently under evaluation as druggable targets, we focused on genes that had this function. Therefore, we selected the three genes included in the functional analyses under the “Histone-lysine N-methyltransferase activity” function, *KMT2D, SETD2* and *SETD1A,* (Figure 1B). Next, we confirmed the presence of these mutations in the different breast cancer subtypes, using data contained at TCGA (Table 1). According to TCGA data, mutations of *KMT2D* were observed in 6% of TNBC and mutations of *SETD2* in 1.2%, confirming the data obtained with METABRIC. However, the presence of mutation in the other breast cancer subtypes was not confirmed or was too low compared to the percentage found in METABRIC. On the other hand, the proportion of *SETD1A* mutations was not confirmed in TCGA for any of the subtypes (Table 1). Next, we aimed to further explore the impact of the mutations of these two genes in patient prognosis, by exploring the effect of their associated transcriptomic signature in breast cancer (All subtypes). *KMT2D* transcriptomic signature was linked with detrimental outcome (HR 0.62 CI: 0.56-0.69; log rank p=0), as well as *SETD1A* (HR 0.66 CI: 0.59-0.74; logrank p=7.6E-14) and *SETD2* (HR 0.69, CI: 0.62-0.77; logrank p=1.8e-11) transcriptomic signatures (Figure 3A).

**Table 1.**
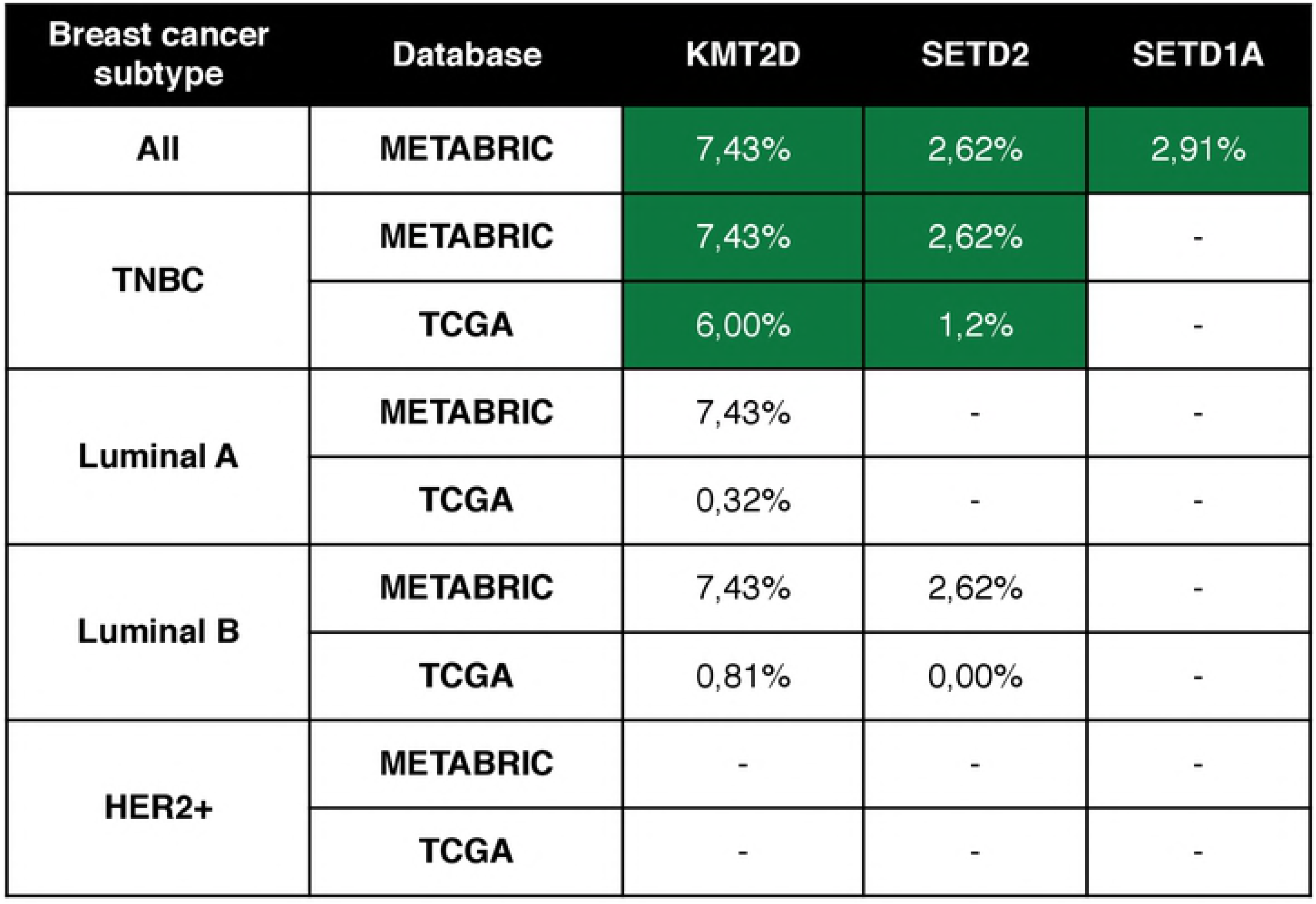
Proportion of mutations in TCGA and METABRIC databases. Proportion of mutations in *KMT2D, SETD2* and *SETD1A* by breast cancer subtype using data from the METABRIC and TCGA studies contained in cBioportal.

As the presence of *KMT2D* and *SETD2* mutations were consistent using both databases (METABRIC and TCGA) in TNBC, we next explored if *KMT2D* and *SETD2* mutational signatures were associated with detrimental prognosis in this specific tumor subtype. Notably, the presence of the associated transcriptomic signatures for both, *KMTD2* and *SETD2,* were associated with poor prognosis (HR 0.58 CI: 0.45-0.74; log rank p=1.9e-05 and HR 0.55 CI: 0.43-0.71; log rank p= 4.2e-0.6; respectively) (Figure 3B).

**Figure 3.**
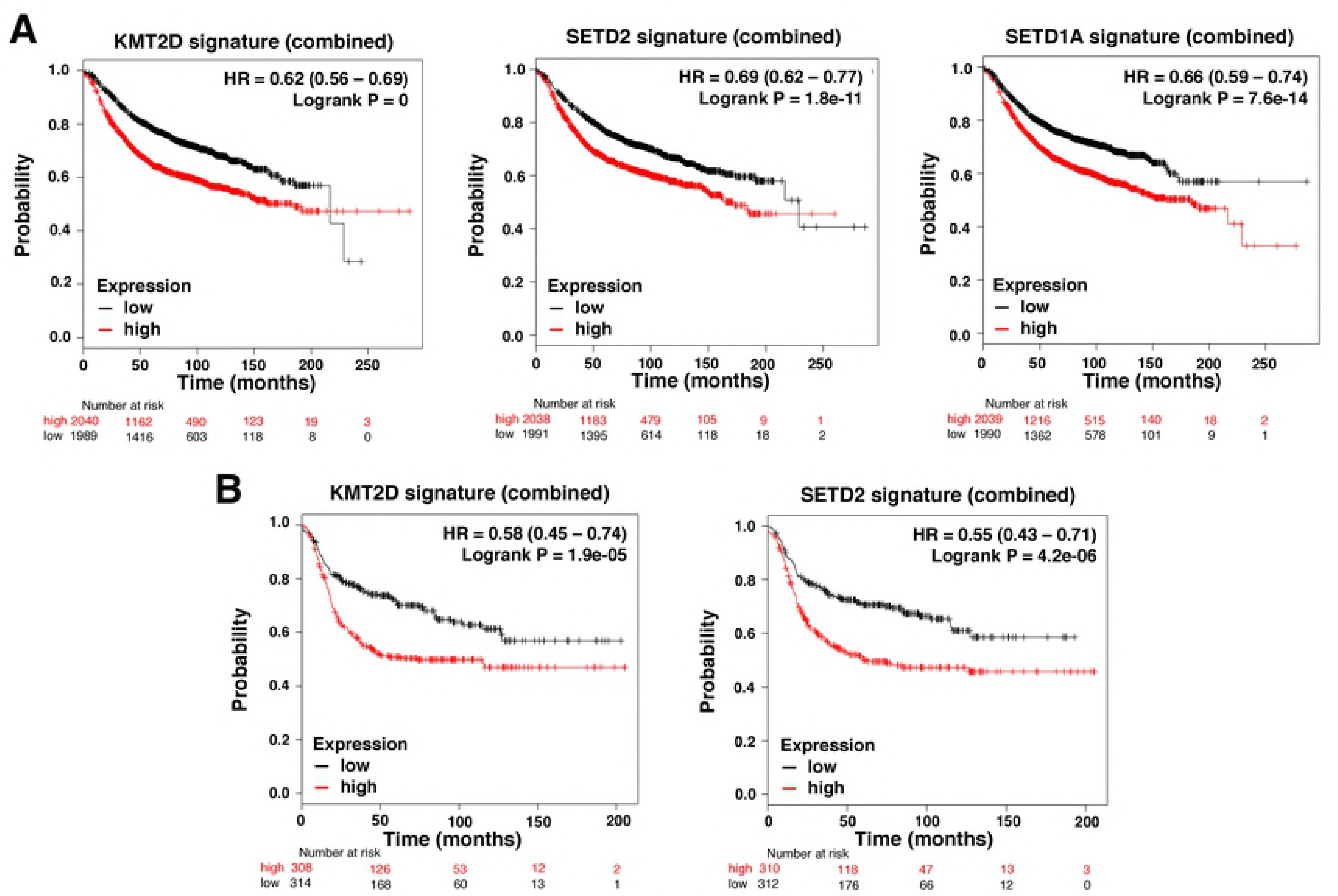
*KMT2D, SETD2* and *SETD1A* mutational signature and clinical outcome. A. Association of *KMT2D, SETD2,* and *SETD1A* mutational signature with patient outcome in all breast tumors. B. Association of *KMT2D* and *SETD2* mutational signatures with prognosis in triple negative breast tumors. The online tool Genotype-2-Outcome was used for both analyses.

From here, we focused on *KMT2D,* as it was the most prevalent mutated gene in both datasets and was strongly associated with poor outcome. KMT2D is a histone methyltransferase that acts as a transcriptional regulator. The complete list of deregulated genes included in the *KMTD2* associated transcriptomic signature is shown in S1 table, and the functions of these genes, determined with the online tool Enrichr, are displayed in figure 4. Most down-regulated genes were included in the cell communication function, followed by tyrosine metabolism or extracellular matrix receptor interaction (S1 Table). Genes which codify for Keratins, *KRT23* or *KRT14,* were among the most relevant genes included in the cell communication function (Figure 4). The most relevant upregulated gene included the GTPase RAC3, that belongs to the RAS family of small GTPases involved in cell proliferation (S1 Table and figure 4) [15].

**Figure 4.**
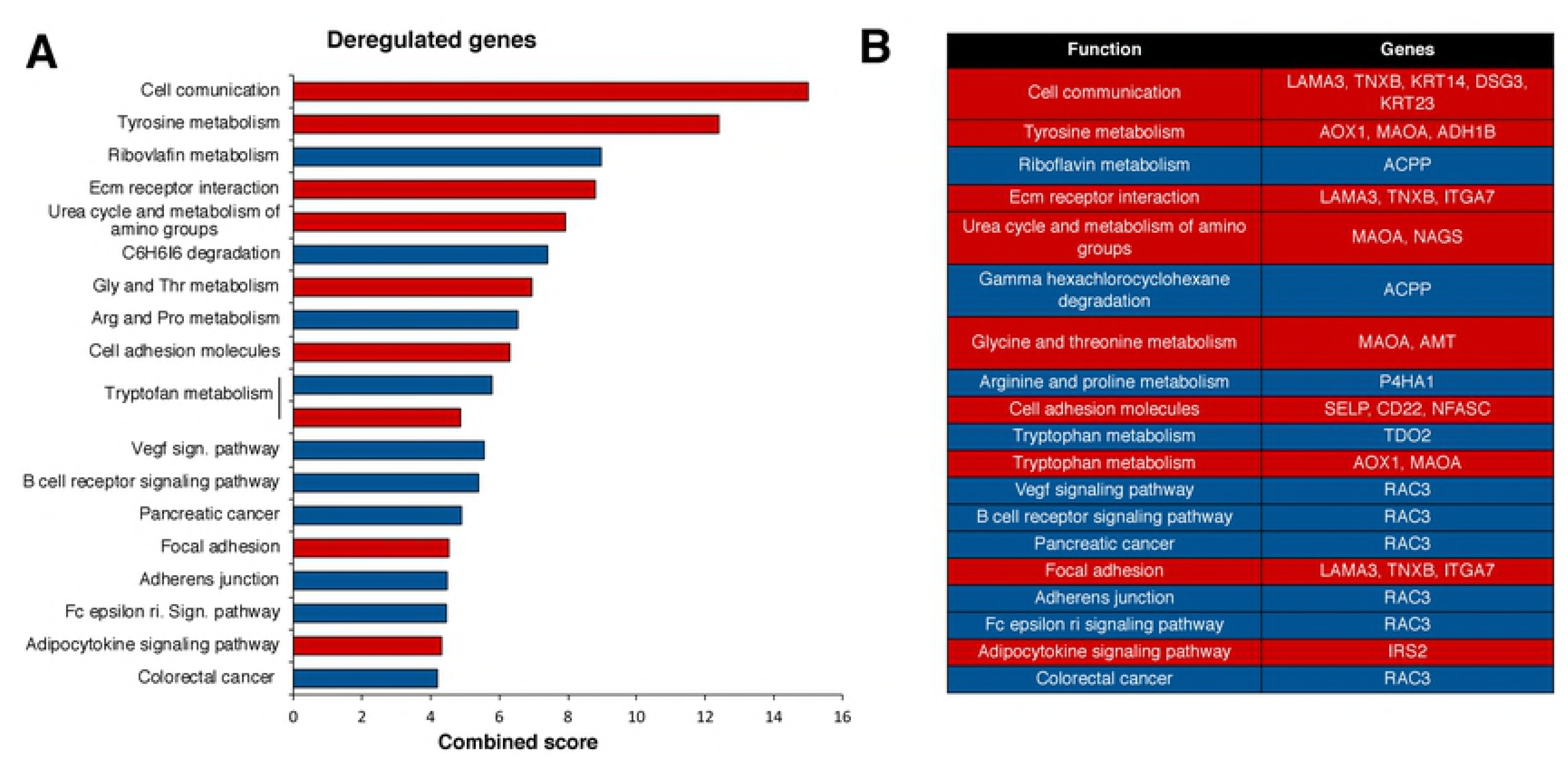
Functional analysis of deregulated genes included in the *KMT2D* mutated signature. A. Percentage of deregulated genes included in the *KMT2D* mutated signature by biological function. Overexpressed genes are displayed in blue and down-expressed genes in red. For functional annotation analysis, the online tool Enrichr was used as described in material and methods. B. Deregulated genes included in each function.

Last, we explored the functional consequences of the mutations present in *KMT2D* in the samples of the METABRIC database. To identify these mutations, we used the online tool cBioportal (Figure 5A). Missense mutations were scattered along the full length of the protein, and were the most abundant molecular alterations, followed by truncating mutations (Figure 5B). The functional impact of all these different mutations, evaluated with three different databases (Mutation Assessor, SIFT and PolyPhen-2), are displayed in Figure 5C. As shown, between 40-55% of *KMT2D* mutations had a functional impact. This indicated that those mutations lead to an abnormal protein, unable to participate in their normal function, mimicking a lack of expression of the gene. To confirm this hypothesis, we decided to explore if a low expression level of this gene could recapitulate the outcome observed at a mutational level, when we explored the effect of mutated *KMT2D.* Using the online tool KMplotter, that links the transcriptional expression of a gene with patient outcome [14], we found that low transcriptomic levels of *KMT2D* were associated with detrimental prognosis (relapse free survival) in all breast tumors (HR 0.64 CI: 0.55-0.79; log rank p=2.4e-08) (Figure 5D), in addition to the triple negative subtype (HR 0.71 CI: 0.551-0.98; log rank p=0.035) (Figure 5E).

**Figure 5.**
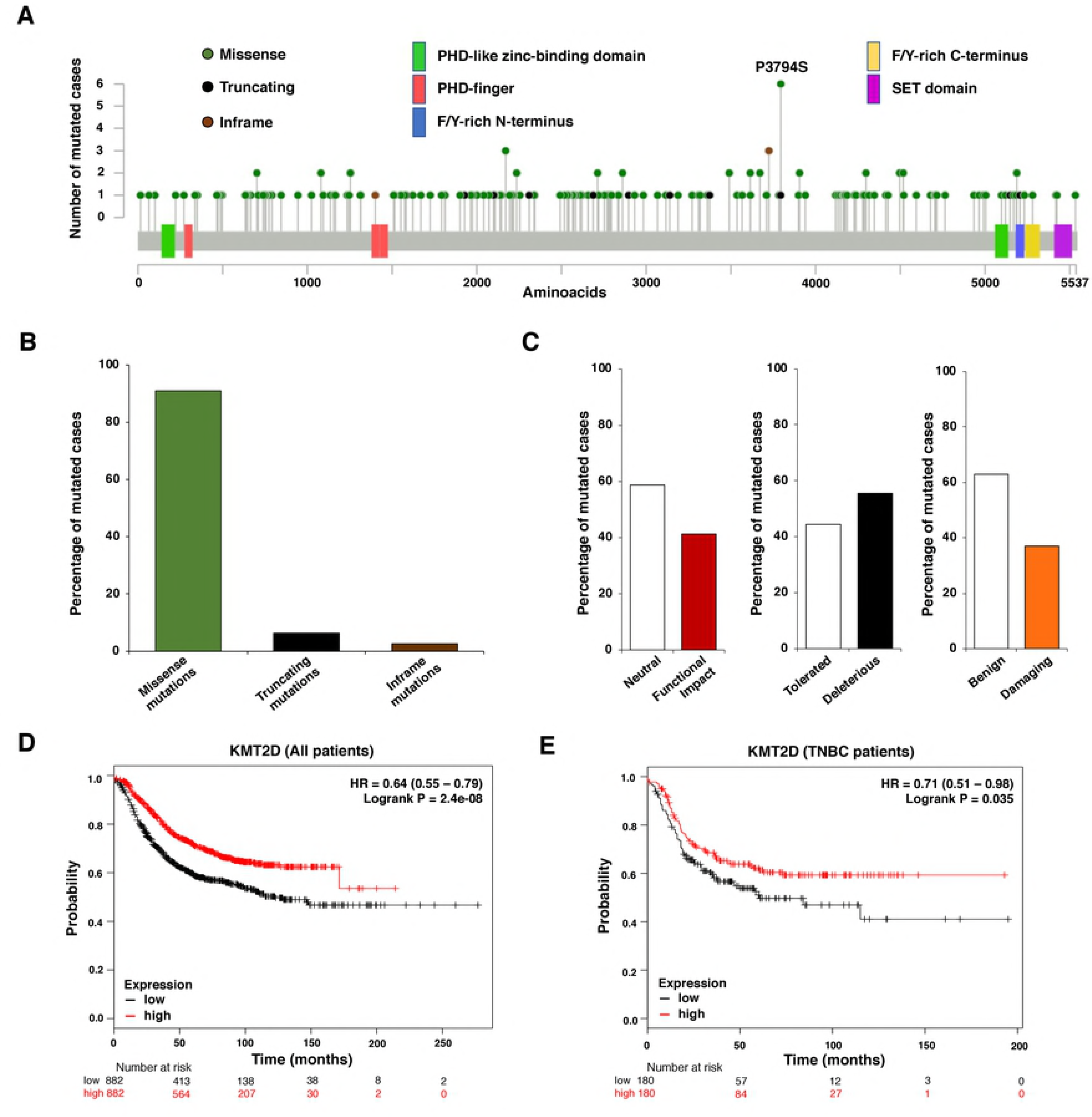
Assessment of mutations at *KMT2D.* A. Diagram showing each aminoacid (aa) which can be found to be mutated in the *KMT2D* gene. B. Type of mutations from the included cases. C. Functional impact of *KMT2D* mutations in the included cases. D. Relapse free survival (RFS) of breast cancer patients based on the transcriptomic expression of *KMT2D.* E. Relapse free survival (RFS) of triple negative breast cancer patients based on the transcriptomic expression of *KMT2D.* KM plotter online tool was used for these prognosis analyses.

## Discussion

In the present article we report the identification of genes that are mutated in breast cancer and associated with detrimental outcome. After functional analysis of the identified genes, we focused on the “Histone-lysine N-methyltransferase activity” function and found that the histone methyltransferase gene *KMT2D* was mutated in around 6% of the TNBCs samples evaluated; in addition to be associated with poor prognosis in this breast cancer subtype.

KMT2D is a histone methyltransferase that methylates the Lys-4 position of the histone H3 [16]. The codified protein belongs to a large protein complex termed ASCOM, which is one of the transcriptional regulators of the estrogen receptor genes [16,17].

*KMT2D* mutations have been associated with the development of different tumors, including small cell lung cancer [16], esophageal squamous cell carcinoma, and large B-cell lymphoma [16]. Although there are many other tumors where mutations in this gene have been described [16,18], neither those mutations have been previously reported in breast cancer, nor their impact on patient outcome has been assessed.

Recent data suggest that KMT2D is involved in the recruitment and activation of relevant breast cancer genes including *FOXA1, PBX1,* and *ER* [17]. As described in the present article and other reports, most of the mutations in *KMT2D* are frameshift and nonsense mutations in the SET and PHD domains, respectively [17]. Most of the described mutations result in the protein loss or in a reduction of the methyltransferase activity [19]. Therefore, this can produce defective enhancer regulation and, subsequently, modifications in the transcription of several genes or increase in genomic instability [8,20]. This is demonstrated in our study by the transcriptomic signature associated with the gene mutation, which will be discussed later, particularly with the upregulation of *RAC3.* Of note, KMT2D displays different effects depending on the cellular context, due to the recruitment of different transcription factors [16].

When evaluating the transcriptomic signature linked to *KMT2D* mutations, we found that *RAC3* was one of the most significantly upregulated transcripts. This transcript codes for a GTPase which belongs to the RAS superfamily of small GTP-binding proteins, and it has been linked with the pathophysiology of many solid tumors, including breast cancer [15,21,22]. In breast cancer *RAC3* regulates invasion and migration participating in the metastatic process[15].

Finally, we confirmed that the expression level of the *KMT2D* gene was associated with clinical outcome in a similar manner as we observed for the presence of the gene mutations, which mostly produce a reduction or loss of protein expression or a decrease in its activity. This result indirectly confirms the robustness of the mutational gene signature in relation to outcome.

In conclusion, in the present work, we identify that the histone methyltransferase gene *KMT2D* is mutated in a number of TNBC patients and associated with detrimental outcome in TNBC. Therefore, modulation of the expression or activity of downstream genes, or *KMT2D* itself, might have relevant consequences from a therapeutic point of view.

## Authors’ Contribution

AO conceived the study and did the original design of the experiments. SMG, MNL, MNC, CNJ and JP searched the data and performed the analysis. AO and EMGM wrote the manuscript. SMG, JP and EMGM prepared the figures. AP, BG, EA, EMGM and AO discussed the data. All authors reviewed, included modification and approved the final version of the manuscript.

## Availability of data and materials

All data generated and/or analyzed during the current study are available from the corresponding author on reasonable request.

## Supplemental material

**S1 Table.** Deregulated genes contained in the transcriptomic signature linked to *KMT2D* mutation. Genes found to be upregulated or downregulated in the *KMT2D* mutational signature. Genotype-2-Outcome database was used for this analysis.

